# NuFold RNA Structure Database: Large-Scale RNA Tertiary Structure Prediction with Confidence Metrics

**DOI:** 10.64898/2026.10.21.760778

**Authors:** David Platt, Yuki Kagaya, Daisuke Kihara

**Affiliations:** Department of Chemical and Biological Engineering, University of Colorado, Boulder, CO, 80310, USA; Department of Biological Sciences, Purdue University, West Lafayette, IN, 47907, USA; Department of Computer Science, Purdue University, West Lafayette, IN, 47907, USA

## Abstract

**Summary:** To bridge the gap between rapid RNA sequence discovery and slow experimental characterization, we present the NuFold RNA Structure Database (https://nufold.kiharalab.org/), a resource providing large-scale, precomputed full-atom RNA tertiary structures. By removing the computational barriers associated with running the method locally, this database makes high-accuracy RNA structure models readily accessible to the biological research community for a wide range of applications.

**Availability and implementation:** All predicted structures and corresponding data are available at https://nufold.kiharalab.org/. The source code for the NuFold program is available at https://github.com/kiharalab/NuFold.

**Supplementary Information:** For more insight into resulting statistics gathered for this project, see https://nufold.kiharalab.org/statistics. For more insight into how to utilize the webserver itself, see https://nufold.kiharalab.org/faq and https://nufold.kiharalab.org/tutorial.

## Introduction

Understanding the three-dimensional structures of non-coding RNAs (ncRNAs) is relevant for elucidating their biological functions and mechanisms, as well as for drug discovery targeting those ncRNAs^1^. While well-characterized ncRNAs, such as tRNAs, rRNAs, and riboswitches, rely on stable tertiary structures to function, for the vast majority of the more than 670,000 Homo sapiens ncRNA sequences reported in RNAcentral^2^, it remains largely unknown whether they adopt well-defined tertiary structures or remain structurally flexible. Among these, only about 500 have experimentally determined structures. Computational approaches provide a promising alternative to explore this uncharted space at scale.

RNA structure prediction is generally more challenging than its protein counterpart, owing to the limited availability of experimentally determined RNA structures and the intrinsic flexibility of RNA molecules. Nevertheless, substantial progress has been made in recent years through the adoption of advanced deep learning approaches. NuFold^3^ is one such method, employing an end-to-end deep neural network architecture similar to AlphaFold2^4^ to directly predict full-atom RNA tertiary structures from input sequences. NuFold uses a flexible nucleobase-center representation that enables torsion-angle control and accurate modeling of ribose ring conformations. Independent benchmarking studies show that NuFold outperforms not only conventional energy-minimization methods but also other recent deep learning-based approaches for RNA monomers as indicated in recent benchmark studies^5,6^. In retrospective benchmark evaluating on CASP15 RNA targets, NuFold achieved the highest global and local structural quality scores among servers for nearly half of the targets, establishing it as a highly accurate and reliable prediction method^3^. Our prediction team using NuFold was ranked third in the CASP16 RNA category^7^.

Given this level of high accuracy, precomputed RNA structure models are expected to be valuable for a wide range of applications, similar to how the AlphaFold Protein Structure Database^8^ has benefited the biological research community. Here, we introduce the NuFold RNA Structure Database, available at https://nufold.kiharalab.org/, which provides large-scale predicted RNA structures generated by NuFold and removes the computational barriers associated with running the method locally.

## Features

The database currently contains 514,124 predicted RNA tertiary structures derived from 48 model organisms. RNA sequences were obtained from RNAcentral, which annotates sequences with organism information and Rfam RNA family classification^9^. For sequence selection, we extracted transcripts containing only standard nucleotides (A, C, G, and U/T) with lengths between 10 to 400 nucleotides. Furthermore, to remove structural prediction artifacts, entries with topological knots detected by RNAspider^10^ were removed. Among all entries, 17.5% (89,909 entries) are classified into 2,243 Rfam families, while the remainder currently lack Rfam annotation in RNAcentral. The sequence lengths range from 10 to 400 nt, with a median of 30 nt (interquartile range: 26-87 nt; mean: 82.5 nt). Across the predicted models, the overall mean pLDDT is 73.2 (median: 72.4), with 62.3% of the entries achieving an average pLDDT > 70. Fig. 1a outlines the computational workflow, where NuFold predicts RNA structures from sequence and predicted secondary structure.

**Figure 1.**
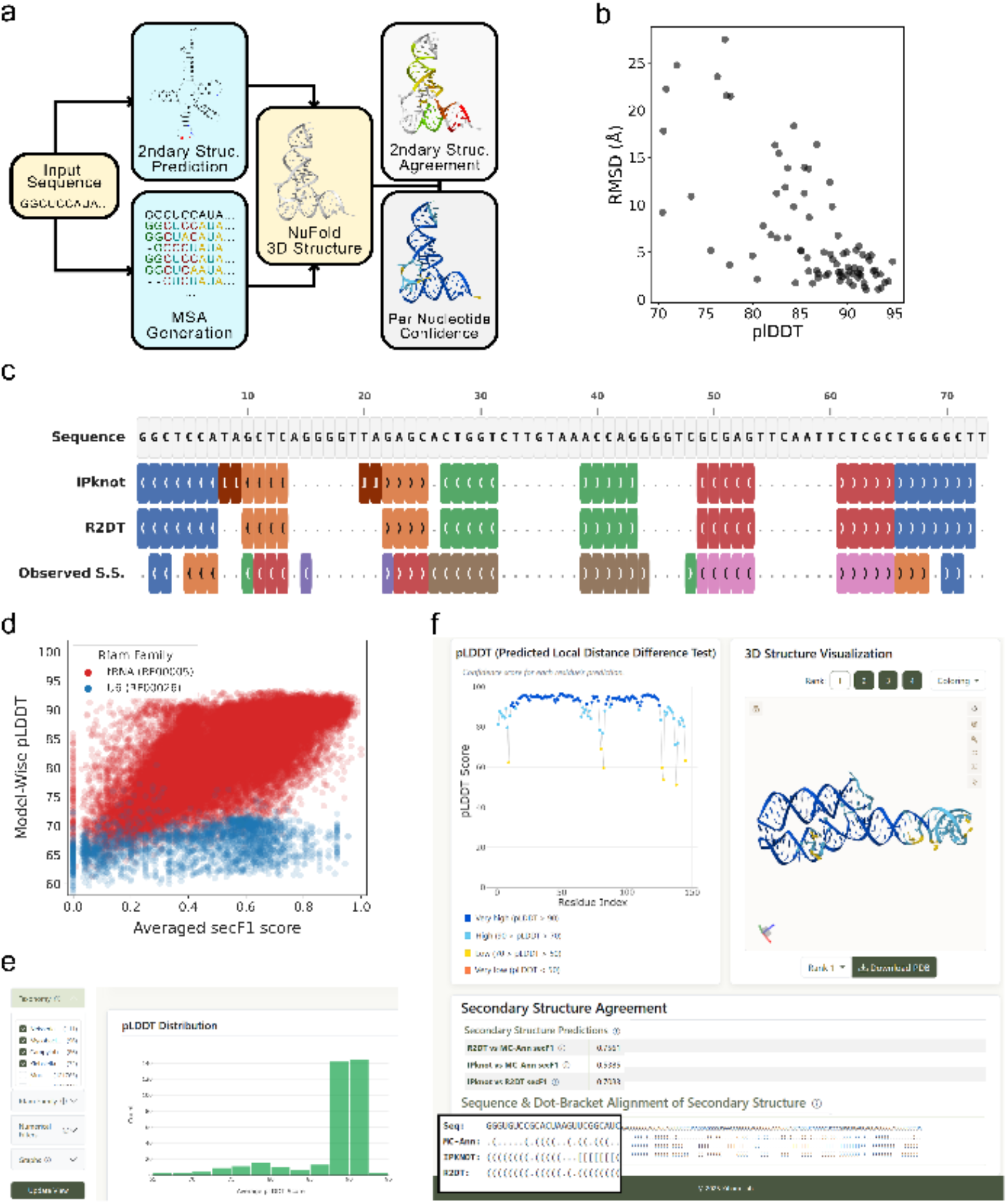
Overview and features of the NuFold RNA Database. (a) Workflow for generating a predicted RNA tertiary structure from an input sequence. The structure model is computed from the input sequence and its predicted secondary structure and is accompanied by two confidence metrics: per-nucleotide confidence (pLDDT) and secondary structure agreement (secF1). The secF1 score evaluates the consistency between secondary structure derived from the model (using MC-Annotate) and that predicted by IPknot or R2DT. (b) Correlation between average pLDDT and root mean square deviation (RMSD) from experimentally determined structures. A total of 84 structures from the validation and test sets reported in the NuFold paper are shown. (c) An example of secondary structure predictions (Homo sapiens tRNA, ID: URS00003516B6_9606). Base pairing clusters are visualized with matching colors. Observed secondary structure was detected by MC-Annotate. (d) Distribution of model-wise pLDDT and average secF1 scores (from IPknot and R2DT) for two RNA families: Homo sapiens tRNA (ID: RF00005; 1432 entries) in red, and Homo sapiens U6 (ID: RF00026; 1314 entries) in blue. (e) An example of the statistics page. Here, users have filtered for 4 specific taxonomies that average high pLDDT plotted via the pLDDT distribution graph. Other graphs can include information about secF1, multiple sequence alignment, and RNA families. (f) An example of NuFold entry. Structure models (4 models) are visualized with multiple choices of color schemes. pLDDT is visualized on the left. Secondary structure agreement is shown in the lower panel.

The NuFold DB places special emphasis on providing model quality assessment, as this is critical information for users. The database reports two complementary accuracy metrics: local geometric confidence (pLDDT) and global secondary structure consistency (secF1 score). Similar to AlphaFold2, NuFold’s pLDDT is a per-nucleotide confidence score ranging from 0 to 100. Benchmarking against experimental structures shows that average pLDDT strongly correlates with RMSD (Fig. 1b). Models with pLDDT > 90 consistently achieve RMSD < 6 Å, indicating reliable overall structural accuracy. While pLDDT effectively captures accuracy in well-structured regions, it penalizes disordered or highly flexible, unpaired segments. To address this limitation, we introduced the secondary structure agreement metric, secF1. It quantifies the agreement between base pairs identified in the RNA structure model using MC-Annotate^11^ and those from reference or predicted secondary structure (Fig. 1c). Specifically, secF1 is computed against R2DT^12^, which serves as the curated reference secondary structure from RNAcentral, as well as IPknot^13^, a secondary structure prediction method used as a part of input to NuFold. Thus, two secF1 scores evaluate consistency with established family consensus templates (RNAcentral) and the fidelity of the 3D model to the secondary structure prediction used in input. Besides serving as an additional metric, secF1 also provides family-specific structural distribution. For most of the RNA families, pLDDT and secF1 have a positive correlation as shown in the example of tRNA family (red, Fig. 1d). On the other hand, some families, for instance, U6 spliceosomal RNAs (blue), often present a range of secF1 values with low pLDDT values. This pattern indicates that the structures possess large, flexible unpaired regions that pLDDT scores poorly.

## Usage

To maximize accessibility, NuFold DB is hosted on an interactive web platform with versatile search and filtering capabilities. Users can navigate directly to specific entries using RNAcentral identifiers (URS IDs) or perform text-based searches using keywords, e.g. organism names and RNA gene names. Search results can be further refined using multi-criteria filters, including taxonomy, Rfam family, sequence length, multiple sequence alignment (MSA) depth, confidence metrics (pLDDT and secF1), and the presence of reference secondary structures. This flexible filtering facilitates targeted research workflows. For example, researchers can swiftly isolate high-confidence tertiary structure candidates within a specific RNA family (such as riboswitches or small regulatory RNAs) by selecting entries with sufficient MSA depth and higher secF1 scores. Furthermore, the filtered results can be sorted by any attribute, and the corresponding list of RNA IDs can be downloaded for batch processing or local record-keeping.

Each RNA entry has a dedicated page displaying structural and quality information (Fig. 1f). For each entry, four predicted models from NuFold are provided, ranked from Model 1 in descending order of average pLDDT scores. This arrangement allows users to inspect and compare alternative predictions alongside the top-ranked model. An integrated Mol* 3D viewer^14^ enables interactive inspection, allowing users to map per-nucleotide pLDDT confidence or assess secondary structure consistency through intuitive color schemes. For large-scale analyses, users can examine database-wide trends via the interactive statistics page (Fig. 1e). Furthermore, complete coordinate archives (PDB files) and associated structural data (including per-nucleotide pLDDT scores, secF1 metrics, predicted secondary structures, and Rfam and organism annotations) can be retrieved via bulk download or programmatically queried through a RESTful public API. This capability facilitates seamless integration into automated bioinformatics pipelines.

## Conclusion

By providing immediate access to over half a million RNA structures with associated validation metrics, the NuFold DB enables accelerated exploration of RNA structures that remain experimentally unresolved. Since its initial release in July 2025, the database has expanded three times, from 93,513 Homo sapiens structures to 514,124 entries across 2,243 Rfam families and 48 model organisms. We will perform periodic updates of the database entries with additional organisms and plan to incorporate other information including multimers, cross-references with experimental structural data, and a partial-sequence match search engine.

## Acknowledgements

This work was supported by the National Institutes of Health [R01GM133840, R35GM158267, and R21AI187928]; and the National Science Foundation [CMMI1825941, MCB1925643, IIS2211598, DMS2151678, DBI2146026, and DBI2003635].

## Author Contributions

D.K. conceived the study. D.P. designed and established the NuFold Database platform. D.P. and Y.K. ran NuFold to generate structure models. Y.K. developed the NuFold program. D.P. analyzed results. D.P. wrote the draft and D.K. critically edited it. All the authors read and approved the manuscript.

## Data Availability

The data underlying this article are available in the NuFold RNA Structure Database at https://nufold.kiharalab.org/. The source code of NuFold used for structure inference is available on GitHub at https://github.com/kiharalab/NuFold.

